# Detecting repeated cancer evolution in human tumours from multi-region sequencing data

**DOI:** 10.1101/156729

**Authors:** Giulio Caravagna, Ylenia Giarratano, Daniele Ramazzotti, Trevor A Graham, Guido Sanguinetti, Andrea Sottoriva

**Affiliations:** School of Informatics, University of Edinburgh, Edinburgh EH8 9AB, UK.; Department of Pathology, Stanford University, California CA 94394, US.; Evolution and Cancer Laboratory, Barts Cancer Institute, Queen Mary University of London, London EC1M 6BQ, UK.; Centre for Evolution and Cancer, The Institute of Cancer Research, London SM2 5NG, UK.

**Author notes:** joint senior authors (Guido Sanguinetti, Andrea Sottoriva). correspondence at and.

## Abstract

Carcinogenesis is an evolutionary process driven by the accumulation of genomic aberrations. Recurrent sequences of genomic changes, both between and within patients, reflect repeated evolution that is valuable for anticipating cancer progression. Multi-region sequencing and phylogenetic analysis allow inference of the partial temporal order of genomic changes within a patient’s tumour. However, the inherent stochasticity of the evolutionary process makes phylogenetic trees from different patients appear very distinct, preventing the robust identification of recurrent evolutionary trajectories. Here we present a novel quantitative method based on a machine learning approach called Transfer Learning (TL) that allows overcoming the stochastic effects of cancer evolution and highlighting hidden recurrences in cancer patient cohorts. When applied to multi-region sequencing datasets from lung, breast and renal cancer (708 samples from 160 patients), our method detected repeated evolutionary trajectories that determine novel patient subgroups, which reproduce in large single-sample cohorts (n=2,641) and have prognostic value. Our method provides a novel patient classification measure that is grounded in the cancer evolution paradigm, and which reveals repeated evolution during tumorigenesis, with implications for our ability to anticipate malignant evolution.

## Introduction

The biggest clinical challenge in oncology is the fact the tumours change over time, progressing from benign to malignant, becoming metastatic, and developing treatment resistance [1, 2]. This occurs through a process of clonal evolution that involve cancer cells and their microenvironment [3]. Intra-tumour heterogeneity, or the genetic and phenotypic variation of cancer cells within the same tumour, is the natural consequence of this evolutionary process and a key factor contributing to the lethal outcome of cancer by providing the substrate of phenotypic variation upon which adaptation can occur [4]. A fundamental question in oncology is therefore: can we predict a cancer’s next evolutionary “step”? Our ability to predict cancer evolution has tremendous implications for clinical care and therefore the question of predictability of evolutionary processes, first posed by evolutionary biologist [5], is central in oncology.

Clonal evolution results from the interplay of random mutations, genetic drift, and non-random selection [6], leading to complex patterns in the data and implying some limits of predictability of cancer evolution due to stochastic forces [7]. However, histopathological staging and molecular markers indicate that, at least in part, tumour evolution is predictable. Moreover, several observations suggest that despite its stochastic nature, microenvironmental, epistatic, and lineage constraints may allow for the prediction of a limited set of subsequent evolutionary moves [2].

Seminal studies have shown how the partial order of somatic aberrations in a patient’s tumour can be determined using multi-region sequencing and phylogenetic analysis, revealing the spatio-temporal dynamics of a single malignancy [8]. However, truncal (clonal) alterations cannot be timed in most cases and phylogenetic trees from different patients often appear very distinct [9, 10, 11, 12, 13, 14]. The underlying variability and complexity of the evolutionary process is such that, with current analysis techniques, we are generally unable to robustly identify recurrent evolutionary trajectories across patients. Characterising repeated evolution in cancer would have severe implications both for understanding the biology of carcinogenesis, and for our ability to stratify patients in the clinic.

Here we exploit the fact that tumours in different patients represent multiple instances of an evolutionary process. This is an opportunity that is missing in classical evolutionary biology, where data come from a unique stream of evolution. To leverage this observation, we devised a novel methodology called REVOLVER (Recurrent EVOLution in cancER) that jointly analyses multi-region sequencing data from patient cohort using a type of machine learning approach called Transfer Learning (TL) [15].

REVOLVER combines the pattern recognition power of machine learning with the knowledge of the underlying biology of tumours to identify clinically relevant recurrent evolutionary trajectories, both between patients and within patients, that are prognostic and can be used to stratify cohorts based on how a patient tumour has evolved (Figure 1).

**Figure 1.**
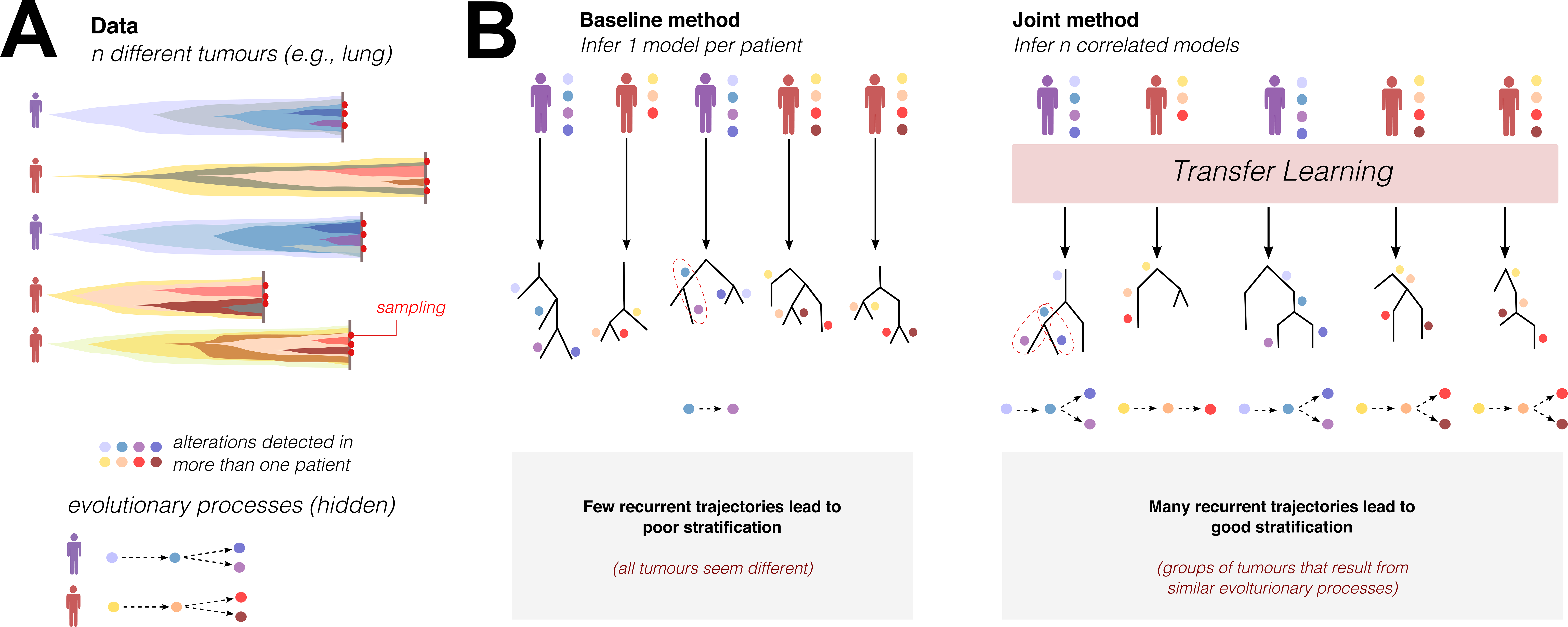
Identifying repeated cancer evolution with Transfer Learning. **A.** Cancer evolution studies often employ multiple sampling of the same tumour and/or sampling of different lesions from the same patient to characterise intra-tumour heterogeneity (ITH). Considering ITH data from *n* patients diagnosed with the same tumour type, we want to find *n* models that describe the evolution of each patient’s malignancy, while at the same time highlighting the presence of recurrent evolutionary trajectories across patients. The aim is to identify and characterise subgroups of tumours that evolved similarly (e.g. group red and group purple), likely driven by similar selective pressures. Due to the underlying stochasticity of cancer evolution, data between patients usually appear very variable and the details of the evolutionary process remain hidden. **B.** The standard approach to this problem is to infer one evolutionary model per patient at a time (e.g. a phylogenetic tree), and then compare the *n* trees. Because in this approach the *n* models are inferred independently and are therefore uncorrelated, as well as due to inherent noise and limited sampling, the statistical signal of recurrent evolutionary trajectories is weak, rendering impossible to confidently identify groups of evolutionary similar tumours. In REVOLVER we use *Transfer Learning* (TL) to infer *n* models jointly with the aim of increasing their structural correlation. We then obtain *n* evolutionary models that besides explaining data in each patients, they highlight the recurrent evolutionary trajectories in the cohort (red dashed circles), thus defining evolutionary subgroups. This results in a stronger statistical signal of recurrent steps in cancer evolution, which we can use to determine the constraints of the evolutionary process in the cohort and potentially inform on tumour progression.

## Results

### REVOLVER: identification of repeated evolution using transfer learning

The standard method to identify recurrent evolutionary trajectories in cancer using multi-region sampling (Figure 1A) is to determine one evolutionary model per patient at a time (e.g. a phylogenetic tree), and then compare the trees. However, since the *n* models are inferred independently and are therefore uncorrelated, the inherent noise in the evolutionary process and in the data renders the statistical signal of recurrent trajectories very weak, making it challenging to reliably identify groups of evolutionary similar tumours (Figure 1B). Our method based on Transfer Learning (TL) instead infers *n* evolutionary models jointly, with the aim of increasing their structural correlation. This allows exploitation of multiple independent observations (single patients) of groups each defined by a (hidden) evolutionary processes (patient evolutionary subgroups) by “transferring” the information to reconcile multiple individual models from the same subgroup (Figure 1B and Supplementary Figure 1). The *n* models still explain the data in each patient, while at the same time highlighting the previously obscure recurrent evolutionary trajectories in the cohort (Figure 1B).

Specifically, our method takes as input intra-tumour heterogeneity data from each patient in the form of a matrix that reports the presence or absence of each alteration in every sample from the tumour. Alterations can be nucleotide substitutions, copy number aberrations (CNA), or any other somatic change. For simplicity, we will refer to any type of alteration as a *mutation*. From the set of data matrices from *n* patients, the method computes *n* trees (*models*) describing the evolutionary trajectories in each tumour. A trajectory is a path *x_1_ → x_2_ → x_3_ →* ••• that connects mutations *x_i_* (Figure 1B and Supplementary Figure 1). An edge *x → y* must satisfy the conditions of temporal precedence (*“x is earlier than y”)* and statistical association (*“x and y are dependent”*). These conditions have been previously used to study cancer evolution from single-sample datasets [16, 17, 18]. For every pair of recurrent mutations *x* and y, our method attempts to find a similar (i.e., correlated) evolutionary trajectory across all patients that harbour *x* and *y*. The aim is to increase the power of the statistical signal of repeated evolutionary trajectories involving recurrent mutations, suggesting constraints to cancer evolution in the cohort.

The genomic features of the different evolutionary subgroups identified using a multi-region profiling dataset (training set) can then be extracted to validate the subgroups in large cross-sectional studies (test set), thus exploiting the combined power of both multi-region and single-sample datasets.

### Recurrent evolutionary trajectories in non-small cell lung cancer

We applied REVOLVER to the TRACERx dataset, the largest multi-region profiling effort to date, comprising the first *n=100* non-small cell lung cancers from the trial [11]. In this cohort, each tumour underwent whole-exome sequencing (*500*x depth) of multiple spatially separated regions *(327* regions in total, median *3* per patient), and a set of putative driver mutations and focal copy number alterations were annotated. In the majority of cases, patients displayed a heterogeneous repertoire of putative driver alterations, often reflecting branched evolution [11].

We examined all putative driver alterations annotated in the original study, and considered recurrent those that appeared in at least 2 patients (Supplementary Figure S4). Tumour suppressor gene inactivation was called in the case of a concurrent mutation and LOH event, or in the case of a homozygous deletion (two-hit inactivation). Oncogenes were separated between mutated and amplified. As reported in the original publication and in previous studies [10], NSCLC is characterised by high intra-tumour heterogeneity at the level of putative driver alterations (Supplementary Figure S5 and S6).

The output of our method are *n=100* correlated models, and several measures of confidence and heterogeneity of the cohort. REVOLVER identified 9 subgroups of recurrent evolutionary trajectories amongst the 100 tumours (Methods; Figure 2A). Subgroup evolutionary trajectories are summarised in Figure 2B. In comparison, clustering the occurrences of driver alterations alone (without TL) does not identify clear subgroups (Supplementary Figure S6). Indeed, previous large-scale single sample studies did not find clinically relevant subgroups using standard approaches [19]. Not surprisingly, a major determinant in the evolution of many lung cancers was TP53 inactivation status. We identified four TP53wt subgroups: G9 characterised by either STK11 or KEAP1 inactivation (statistically indistinguishable with this cohort size), G7 with EGFR mutation, G4 with KRAS mutation alone and G5 with KRAS mutation followed by MGA mutation. Interestingly, the TP53 inactivated subgroups displayed convergence through alternative evolutionary trajectories, one where TP53 was an early, possibly initiating lesion (G1, G2 and G3), and one where it was a late subclonal alteration (G6). Subgroup G3 was characterised by an early TP53 lesion alone that could then branch into subclonal CDKN2A inactivation (mostly due to homozygous deletions) or PIK3CA mutation (G1) or KRAS mutation (G2). Strikingly, subgroup G6 converged to subclonal TP53 inactivation after one of four alternative initial events: MGA, PIK3CA or NFE2L2 mutation, or CDKN2A inactivation – again statistically indistinguishable given the size of the subgroup. The same alternative convergence was observed for PIK3CA mutations in G6 versus G1. This suggests a remarkable level of evolutionary convergence onto recurrent lesions across patients. An example of a patient belonging to an evolutionary subgroup is reported in Figure 2C, although additional sequences of alterations could be present in individual patients, these are not sufficiently recurrent within the cohort. We also identified a subgroup G8 with unknown driver events. We note that we cannot exclude further branching in the evolutionary trajectories as a consequence of alterations that are not recorded in this dataset, such as epigenetic changes (e.g. in G8).

**Figure 2.**
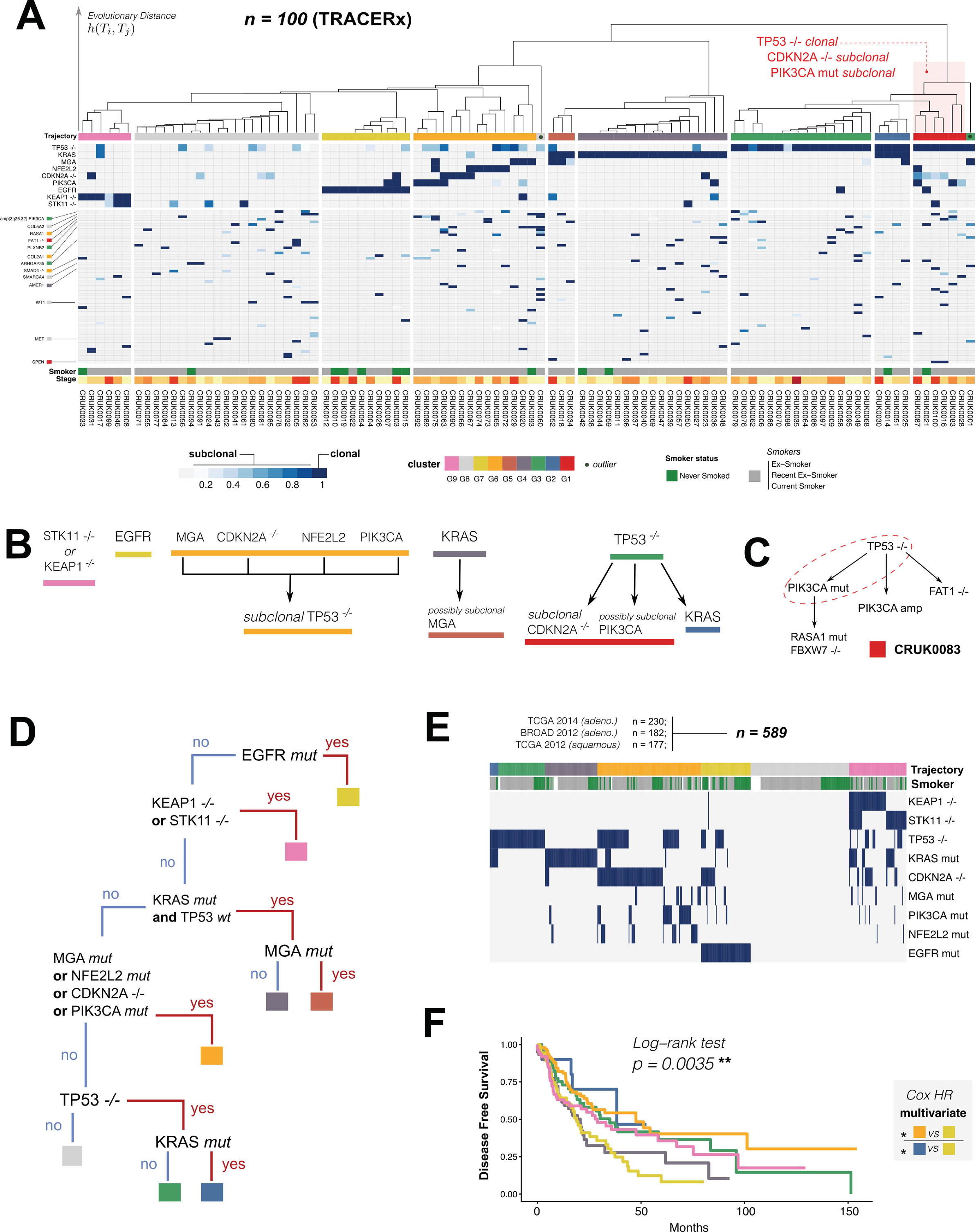
Recurrent evolutionary trajectories in lung cancer. **A.** Application of REVOLVER to *n=100* lung cancers sequenced within the TRACERx project [11] allowed stratification of patients in 9 evolutionary subgroups. Each column is a patient; each row a genomic event: inactivation of a tumour suppressor, activation of an oncogene, or mutation of a gene annotated as driver in [11]; the blue intensity shows if the mutation is clonal or subclonal in the patient. To obtain these groups we processed the putative driver genes in TRACERx, and considered as recurrent all alterations observed in at least 2 patients. The output of our method are *n=100* correlated models for the cohort, which we stratify using a distance metric *h* (in subgroup G1, the characteristic events are annotated). Covariates such as smoker status and stage are also reported on the bottom of the heatmap. **B.** We report the most likely trajectories that are recurrent in each group, and whether they involve clonal or subclonal alterations. Interestingly, the TP53 inactivated subgroups displayed convergence through alternative evolutionary trajectories, one where TP53 was an early, possibly initiating lesion (G1, G2 and G3), and one where it was a late subclonal alteration (G6). Similar patterns were identified for PIK3CA (G6 versus G1). This suggests a remarkable level of evolutionary convergence onto recurrent lesions across patients. **C.** Example of a patient belonging to subgroup G1; although further branching evolution could occur in an individual patient, the subgroup analysis highlights recurrent branching in the cohort, identifying trajectories that repeat in multiple patients. **D.** With the 9 groups, we can generate a decision tree classifier from the genomic events characterizing each group, and use it to classify single-sample tumours. The low-resolution of single-sample data, however, does not allow to disentangle, for instance, clonal from subclonal TP53 inactivation, and hence G1 from G6 tumours. Here we prioritise the larger group G6. **E.** Our groups validate against a cross-sectional test set of *n=589* tumours from 2 TCGA and 1 Broad Institute projects [20, 21, 22], and also show significant enrichment for smoking and never-smoking status. **F.** Kaplan Meier analysis identified significant differences in disease-free survival amongst the different subgroups (log-rank, *p=0.0035*). Results from multivariate analysis showed significance in G6 versus G7 (HR 1.94, CI (1.05; 3.6), *p=0.03)* and G2 versus G7 (HR 3.13, CI (1.02; 9.61), *p=0.04*).

From the signatures of each group we derived a decision tree (Figure 2D) to classify single-sample dataset and validate our subgroups. However, since we could not accurately distinguish subclonal from clonal TP53^-/-^ or PIK3CA mutations from publically available single-sample processed data, we could not distinguish G1 from G6 in those large datasets and we chose to prioritise the larger group G6. With the decision tree constructed from TRACERx’s multi-region sequencing data we stratified *n=589* tumours from two TCGA and one Broad Institute studies [20, 21, 22] (Figure 2E).

The evolutionary subgroups replicate in these large datasets, confirming the robustness of our stratification. Remarkably, Kaplan-Meier analysis identified significant differences in disease-free survival between the groups (Figure 2F, *log-rank p=0.0035*). Multivariate Cox Proportional Hazards (corrected for age, stage, histology, packs per year and smoker status) confirmed significance for several groups, notably G6 versus G7 (HR 1.94, Confidence Interval (1.05; 3.6) *p=0.034)* and G2 versus G7 (HR 3.13, Confidence Interval (1.02; 9.61) *p=0.046)* (Figure 2F and Supplementary Figure S7). Furthermore, non-smoker status was significantly enriched in EGFR-triggered tumours (G7, *Fisher p<0.001),* while smoker status was enriched in tumours with inactivated KEAP1 or STK11 (G9, *Fisher p=0.015).*

### Recurrent evolutionary trajectories in breast cancer

We applied REVOLVER to a cohort of *n=50* primary breast cancers where multi-region whole-genome and targeted bulk sequencing was available (303 samples in total) [14]. In each sample, a panel of mutations and CNAs (cytoband-level and whole-arm) in breast cancer putative driver genes were annotated (Supplementary Figure S8). We processed all annotated mutations and CNAs, and considered recurrent those appearing in at least 2 patients, as we did for the TRACERx cohort. REVOLVER stratified this cohort in 5 evolutionary groups (Figure 3A, Supplementary Figure S9 and S10). As we found in lung cancer and as reported in the original study [14], recurrent evolutionary trajectories could not be identified by analysing the occurrences of driver alterations alone (Supplementary Figure S9). From recurrent trajectories (Figure 3B) we computed a decision tree (Figure 3D), which shows that TP53 is a key node in the evolutionary history of breast cancer, most likely also in this case initiating genomic instability. Then the two most recurrent chromosomal alterations in breast cancer, 17p loss and 8q gain define branching but convergent trajectories for both TP53 mutant and wt cancers (G2 and G3). Moreover, cancer cells with TP53 mutation, 17p loss and 8q gain can further acquire MYC amplification (G4); an example patient from this subgroup is shown in Figure 3C.

**Figure 3.**
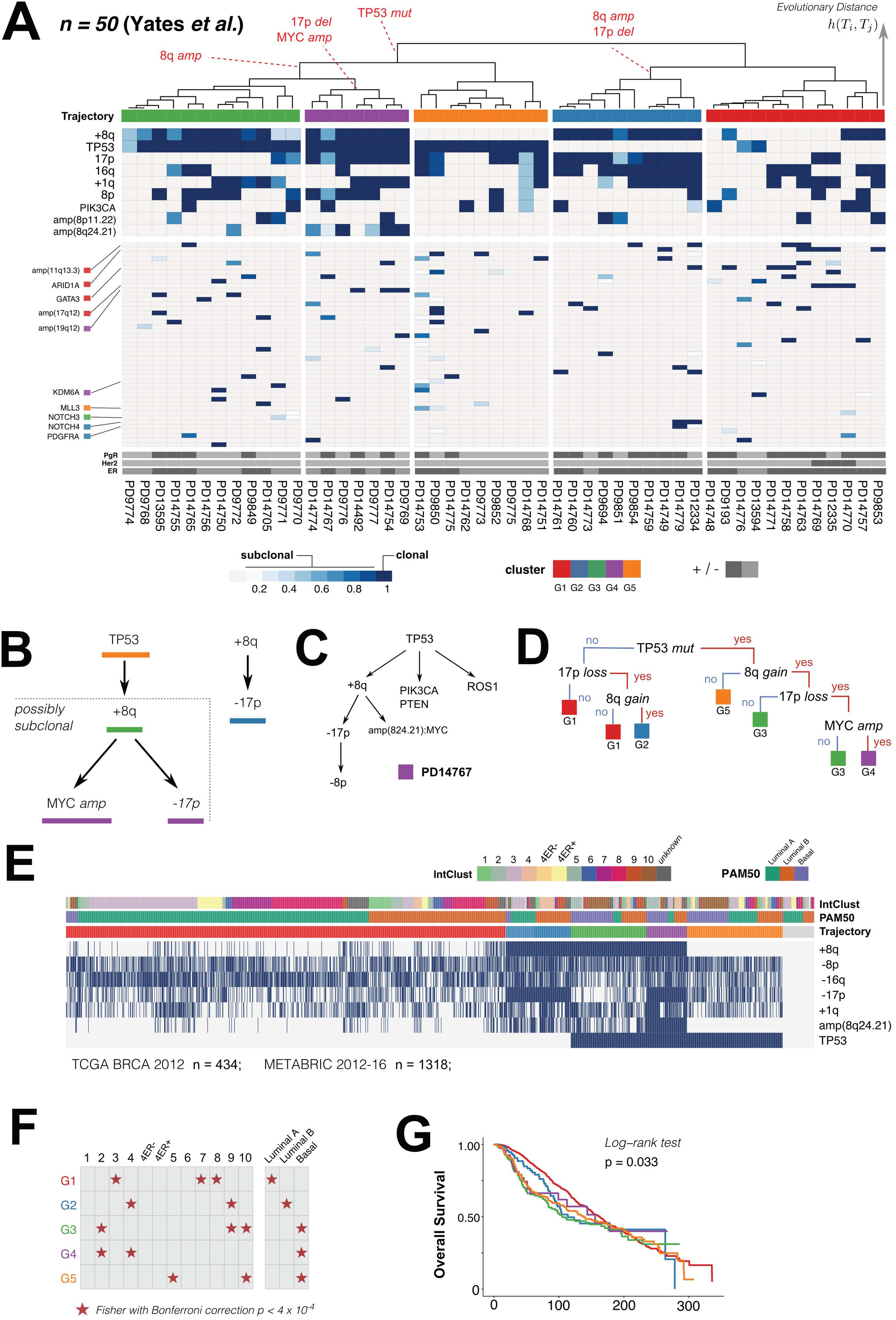
Recurrent evolutionary trajectories in breast cancer. **A.** Application of REVOLVER to analyse *n=50* breast cancer patients [14] results in *5* groups of evolutionary trajectories. To analyse this cohort, we process all putative driver mutations/ CNAs annotated in [14]; we consider recurrent those detected in at least 2 patients. **B.** A decision tree classifier consistent with the identified groups. **C.** Example of a patient belonging to subgroup G4. D. We validate our groups on *n=1,752* single-sample breast tumours collected by TCGA and METABRIC projects [23,24], which we stratify by using the decision tree. **D.** The groups show significant (Fisher test, *p<0.05)* enrichment for IntClust and PAM50 classifications, whose phenotypes associated to different groups. For instance, IntClust 5 associated to G5, and IntClust 10 to G3 and G5; and Basal tumours to TP53 mutated groups (G3, G4 and G5). **E.** Kaplan Meier plots and log-rank tests show significant survival outcome among the identified groups (log-rank test, *p<0.05).*

We again used the decision tree from the multi-region dataset to build a classifier and applied it to *n=1,752* single-sample breast cancer cases from the METABRIC *(n=1,318)* and BRCA TCGA *(n=434)* studies [23, 24] and found that our evolutionary subgroups replicated (Figure 3E). The evolutionary subgroups were enriched for specific breast cancer subtypes from the IntClust (based on both transcriptomic and copy number) and PAM50 (transcriptomic alone) classifications (Figure 3F), for example IntClust 5 and 10 were associated with G3 and G5 tumours, while Luminal A subtypes were associated with G1 and Luminal B with G2. The fact that some evolutionary trajectories were associated with multiple IntClust/PAM50 subtypes suggests convergent evolution at the level of phenotype (transcriptome), for example IntClust 2 was associated with both G3 and G4 whereas Basals were associated with G3, G4 and G5. Overall survival analysis was also significantly different between groups (Figure 3E, log-rank *p=0.033).*

This analysis show that, as in the case of lung cancer, REVOLVER can recapitulate known biology and provide an effective tool for a prognostically valuable stratification of breast cancer patients based on how their tumours evolved.

### Recurrent evolutionary trajectories in renal carcinoma

We also analysed mutations annotated in *n=10* clear cell renal cell carcinomas collected in [9]. Despite the small number of patients, the high number of samples per tumour allowed the identification of three subgroups (Figure 4A). In this cohort, all patients were characterised by VHL mutation. Subgroup G1 was identifiable by VHL mutation alone, G2 by VHL and PBRM1 together, and G3 by VHL and BAP1 together (Figure 4B). Loss of PBRM1 expression is associated with renal cell carcinoma progression [25], and known expression subtypes for clear cell renal cell carcinomas are associated with BAP1 and PBRM1 [26]. Moreover, BAP1 and PBRM1 mutations are largely mutually exclusive and related to divergent clinical behaviours, with BAP1-mutant tumours associated with worst prognosis [27]. We used the decision tree to validate these subgroups in a large single-sample cross-sectional TCGA cohort of *n=300* kidney tumours [28] and found that the subgroups validated (Figure 4C), although an additional subgroup with no evidence of VHL alteration was present in this larger cohort. Interestingly, survival analysis confirmed the worse prognosis of the BAP1 mutant cases (Figure 4D). The unknown group, containing a mixture of mutually exclusive PBRM1 and BAP1 mutant cancers also showed worse prognosis with respect to G1 and G2, likely driven by the BAP1 cases in the group. Multivariate analysis of the Hazard Ratios confirms the significance in survival risk due to the presence of BAP1/PBRM1 in the unknown group, against their absence in G1’s (VHL-mutated tumours; HR 2.05, Confidence Interval (1.14; 3.67) *p=0.015*).

**Figure 4.**
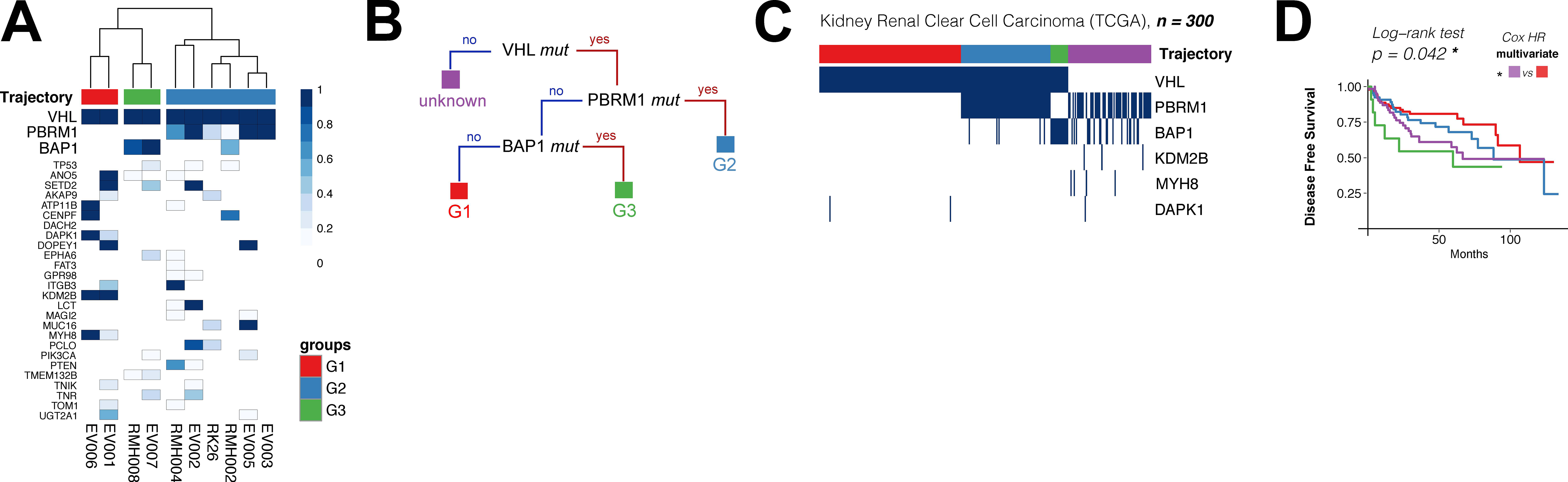
Recurrent evolutionary trajectories in renal cancer. **A.** We analyzed with REVOLVER mutations in *n=10* clear cell renal cell carcinomas (CCRCC) originally published by Gerlinger et al. [9] and identified 3 subgroups. **B.** The decision tree shows the trajectories of the three groups as well as an additional unknown group with no VHL inactivation, which was not present in the multi-region cohort, but it is present in other larger studies. **C.** The validation was carried out using *n=300* samples available in the TCGA kidney cancer cohort [28]. **D.** Survival analysis shows significant difference in the outcome of the groups, both with log-rank test and multivariate Cox regression analysis, highlighting the worse prognosis of the BAP1 mutated tumours.

## Discussion

Detecting repeated evolution in cancer is critical for the implementation of evolutionary approaches to disease management. Stratifying patients based on their recurrent evolutionary patterns facilitates the prediction of the future steps in the evolutionary process of their malignancy, thus potentially allowing taking optimal and personalised clinical decisions.

Although the application of ‘artificial intelligence’ algorithms based on machine learning methods to biomedical datasets is becoming popular [29], the use of these methods as ‘black boxes’ to mine cancer genomic data is unlikely to be successful if not combined with the extensive clinical and biological knowledge we have of human malignancies.

Here we have combined a novel approach based on Transfer Learning, a relatively new set of machine learning methods, with the current knowledge of tumour biology and evolution, to detect the hidden signal of recurrent evolution within multiple tumour types. We have demonstrated how apparently hidden evolutionary trajectories can be identified using this method. Our approach also helps to reconcile multi-region sequencing data with large single-sample cohorts by combining different data types and extracting more information on the evolutionary process from both strategies concurrently. As our method is flexible in terms of input data, the stratification power could be further increased with larger datasets, more accurate technologies (e.g. single-cell sequencing) and longitudinal data obtained, for instance, from ctDNA. The evolutionary subgroups we identified have prognostic value, demonstrating the likely clinical value of stratifying tumours based upon their evolutionary trajectories.

### Authors’ contributions

GC, GS and AS designed the research. GC and GS defined the method. GC and YG implemented it. GC, YG and DR analysed the data. All authors interpreted the results, drafted and approved the manuscript.

